# Higher amplitudes of visual networks are associated with trait-but not state-depression

**DOI:** 10.1101/2024.03.25.584801

**Authors:** Wei Zhang, Rosie Dutt, Daphne Lew, Deanna M. Barch, Janine Bijsterbosch

**Author notes:** Corresponding authors: Wei Zhang; Janine Bijsterbosch.

## Abstract

Despite depression being a leading cause of global disability, neuroimaging studies have struggled to identify replicable neural correlates of depression or explain limited variance. This challenge may, in part, stem from the intertwined state (current symptoms; variable) and trait (general propensity; stable) experiences of depression.

Here, we sought to disentangle state from trait experiences of depression by leveraging a longitudinal cohort and stratifying individuals into four groups: those in remission (‘trait depression group’), those with large longitudinal severity changes in depression symptomatology (‘state depression group’), and their respective matched control groups (total analytic n=1,030). We hypothesized that spatial network organization would be linked to trait depression due to its temporal stability, whereas functional connectivity between networks would be more sensitive to state-dependent depression symptoms due to its capacity to fluctuate.

We identified 15 large-scale probabilistic functional networks from resting-state fMRI data and performed group comparisons on the amplitude, connectivity, and spatial overlap between these networks, using matched control participants as reference. Our findings revealed higher amplitude in visual networks for the trait depression group at the time of remission, in contrast to controls. This observation may suggest altered visual processing in individuals predisposed to developing depression over time. No significant group differences were observed in any other network measures for the trait-control comparison, nor in any measures for the state-control comparison. These results underscore the overlooked contribution of visual networks to the psychopathology of depression and provide evidence for distinct neural correlates between state and trait experiences of depression.

## Introduction

Depression is a global health challenge, emerging as the foremost cause of disability and affecting more than 300 million individuals worldwide (Friedrich, 2017). While there is widespread acknowledgment of depression as a disorder associated with dysfunctions of large-scale brain networks (Williams, 2016), existing research has been hindered by inconsistencies (Greene et al., 2022; Tozzi et al., 2020; Xia et al., 2019), lack of reproducibility (Kennis et al., 2020; Saberi et al., 2022), and, at best, the ability to account for only a modest proportion of variance (Dutt et al., 2022; Schmaal, 2022; Winter et al., 2022). This prevailing uncertainty creates a substantial gap in our comprehension of the neural basis and potential etiology of this pervasive mental health condition. Addressing this knowledge deficit is important, as a deeper understanding of depression’s etiological underpinnings can improve diagnostic, treatment, and prevention strategies.

At the functional neurocircuitry level, investigations using resting-state fMRI have revealed key insights into the differences between individuals with major depressive disorder (MDD) and healthy controls (Kaiser et al., 2015; Mulders et al., 2015). Importantly, previous review and meta-analysis studies on depression indicate hyperconnectivity within the default mode network (DMN) and hypoconnectivity within the central executive network (CEN) (Kaiser et al., 2015). However, recent meta- and mega-analyses present contradictory findings, reporting comparable connectivity patterns of the DMN and CEN between patients and controls (Javaheripour et al., 2021), or reduced connectivity in patients (C.-G. Yan et al., 2019). Despite being less explored in previous neuroimaging studies on depression, disruptions in the functional connectivity of sensory and motor networks have also been implicated in MDD, albeit with similar inconsistency in directions of these connectivity disruptions both within sensory and motor networks or between these and other brain networks (L. Kang et al., 2018; Lu et al., 2020; Ray et al., 2021; Wu et al., 2023; Wüthrich et al., 2023; Zeng et al., 2012; Zhu et al., 2021).

The varying findings across studies can be attributed, at least in part, to the diverse study designs employed. These include comparisons between patient and control groups (Flint et al., 2021; Winter et al., 2022), regressions against depression severity scores (Oathes et al., 2015; Yoshida et al., 2017) or personality traits like neuroticism (Braund et al., 2022; Fournier et al., 2017; Steffens et al., 2017), and investigations into heterogeneity (Dinga et al., 2019; Drysdale et al., 2017; Hannon et al., 2022; Wen et al., 2022; Yang et al., 2021). Moreover, the mixture of state depression (current symptom severity) and trait depression (long-term propensity for depression) further complicates study designs. While state and trait depression are often viewed as closely related and sometimes used interchangeably, biomarkers that differentiate between state and trait depression can serve distinct purposes in a clinical setting. For instance, trait biomarkers are instrumental in identifying individuals at risk, while state biomarkers help gauge treatment effectiveness and track patients’ progress over time. This differentiation aligns with established categories of biomarkers outlined by the FDA-NIH Biomarker Working Group (Group, 2021). The failure to differentiate between state and trait depression likely contributes to inconsistencies in findings across the literature. Therefore, elucidating the distinct neural correlates of state versus trait depression is poised to not only enhance the utility of biomarkers but also offer clarity regarding the underlying neural mechanisms of the disorder.

To date, few studies have specifically examined state and/or trait depression. These investigations often involve comparisons between participants in remission (trait) and those experiencing a current episode of depression (state) (Admon et al., 2015; Ming et al., 2017), or employ longitudinal designs to assess changes pre- and post-pharmacological interventions (state) (Delaveau et al., 2011). Unfortunately, however, these studies have yielded mixed results. In one of those studies, the ventromedial prefrontal cortex and precuneus were suggested to signify trait markers of depression (Ming et al., 2017), while another study indicated altered activity in the same regions following antidepressant drug treatment, implying state-dependent changes (Delaveau et al., 2011). Consequently, the potentially dissociable neural correlates of state versus trait depression remain inadequately understood.

Leveraging the UK Biobank (UKB) data, the present study aimed to identify potentially dissociable resting-state functional correlates of state and trait depression. Inspired by previous work (Admon et al., 2015; Delaveau et al., 2011; Ming et al., 2017), we employed longitudinal data of depression severity to differentiate state and trait experiences. This approach resulted in indications of a high-level propensity to depression during remission (trait) and substantial fluctuations in symptom severities between two time points (state), respectively, effectively dissociating state from trait depression. We further applied a state-of-the-art data-driven decomposition method to identify large-scale brain networks from the resting-state fMRI data and estimated the amplitude (i.e., network strength), connectivity (i.e., between-network temporal correlations also known as network matrices), and spatial overlap (i.e., shared regions between network spatial maps) of these brain networks. Building upon literature indicating dynamic alterations in functional connectivity and more enduring spatial configurations of brain networks (Harrison et al., 2020), we hypothesized that longitudinal changes in functional connectivity of large-scale brain networks would be associated with changes in depression symptom severity (i.e., state depression), whereas spatial organizations of those networks would be indicative of trait depression experiences.

## Methods

The group sampling and statistical analysis plans described below were pre-registered at the Open Science Framework (Zhang & Bijsterbosch, 2023).

### 1. Participants

Out of N=5,215 longitudinal UKB participants, N=4,595 had complete longitudinal neuroimaging and depression data. To discern state and trait depression, we created two corresponding groups and matched controls. The state depression group (N=311) was based on change scores in depression severity between two assessments (i.e., scan 1 and scan 2), while the trait depression group (N=265) had high baseline (i.e., high propensity) but low current symptom severity (i.e., remission at scan 1). Their respectively matched controls (N=311 and N=265) had consistently minimal severity scores across all time points (see detailed group definitions in *section 2*). The resulting sample (N=1,030) included n=33 overlaps between the state and trait groups and n=89 overlaps between the two matched control groups (see study sample flowchart in ***Figure 1A***). As we performed statistical analyses separately for state-control and trait-control comparisons, these overlapping participants were included in each group’s respective comparison analysis. Demographics are summarized in ***Table 1***.

**Table 1.** Demographics of Matched Depression and Control Groups.

|  | Matched State-Control Groups |  |  | Matched Trait-Control Groups |  |  |
| --- | --- | --- | --- | --- | --- | --- |
|  | State<br>(n=311) | Control<br>(n=311) | Stats<sup>a</sup> | Trait<br>(n=265) | Control<br>(n=265) | Stats<sup>a</sup> |
| N<sub>Female</sub> (%) | 196 (63) | 217 (70) | 3.18 | 167 (63) | 179 (68) | 1.20 |
| Mean<sub>age-baseline</sub> (SD) | 51.21 (6.79) | 51.38 (7.29) | 0.32 | 52.19 (6.49) | 52.06 (7.26) | -0.23 |
| Mean<sub>age-scan1</sub> (SD) | 59.95 (6.58) | 60.22 (7.26) | 0.49 | 61.01 (6.55) | 60.81 (7.26) | -0.36 |
| Mean<sub>age-scan2</sub> (SD) | 62.61 (6.39) | 62.83 (4.77) | 0.41 | 63.43 (4.40) | 63.43 (7.04) | -0.53 |
| Mean<sub>RDS-baseline</sub> (SD) | 6.39 (2.48) | 4.77 (0.42) | -12.38** | 7.82 (1.33) | 4.40 (0.49) | -39.00** |
| Mean<sub>RDS-scan1</sub> (SD) | 7.14 (2.73) | 4.31 (0.46) | -18.09** | 4.57 (0.50) | 4.56 (0.50) | -0.71 |
| Mean<sub>RDS-scan2</sub> (SD) | 7.30 (2.91) | 4.31 (0.46) | -17.97** | 5.26 (1.48) | 5.26 (1.48) | -7.79** |
| Mean<sub>interval</sub> (SD)<sup>b</sup> | 949 (360) | 973 (423) | 1.09 | 3221 (492) | 3214 (514) | -1.15 |
| N<sub>Antidepressants</sub><sup>c</sup> | 79 | 0 | NA | 38 | 0 | NA |

N = sample size, SD = standard deviation, RDS = Recent Depressive Symptom scale, NA = not applicable.
<sup>a</sup> Chi-square test was conducted to examine between-group difference in sex, and unpaired t-tests to examine numerical variables.
<sup>b</sup> Intervals (in days) between baseline assessment and scan1 for state, between scan1 and scan2 for trait.
<sup>c</sup> Number of participants on antidepressants in each group (see *Supplemental Materials* for the full list of medications). By definition, no control participants had antidepressants.
\*\*p<0.0001

**Figure 1.**
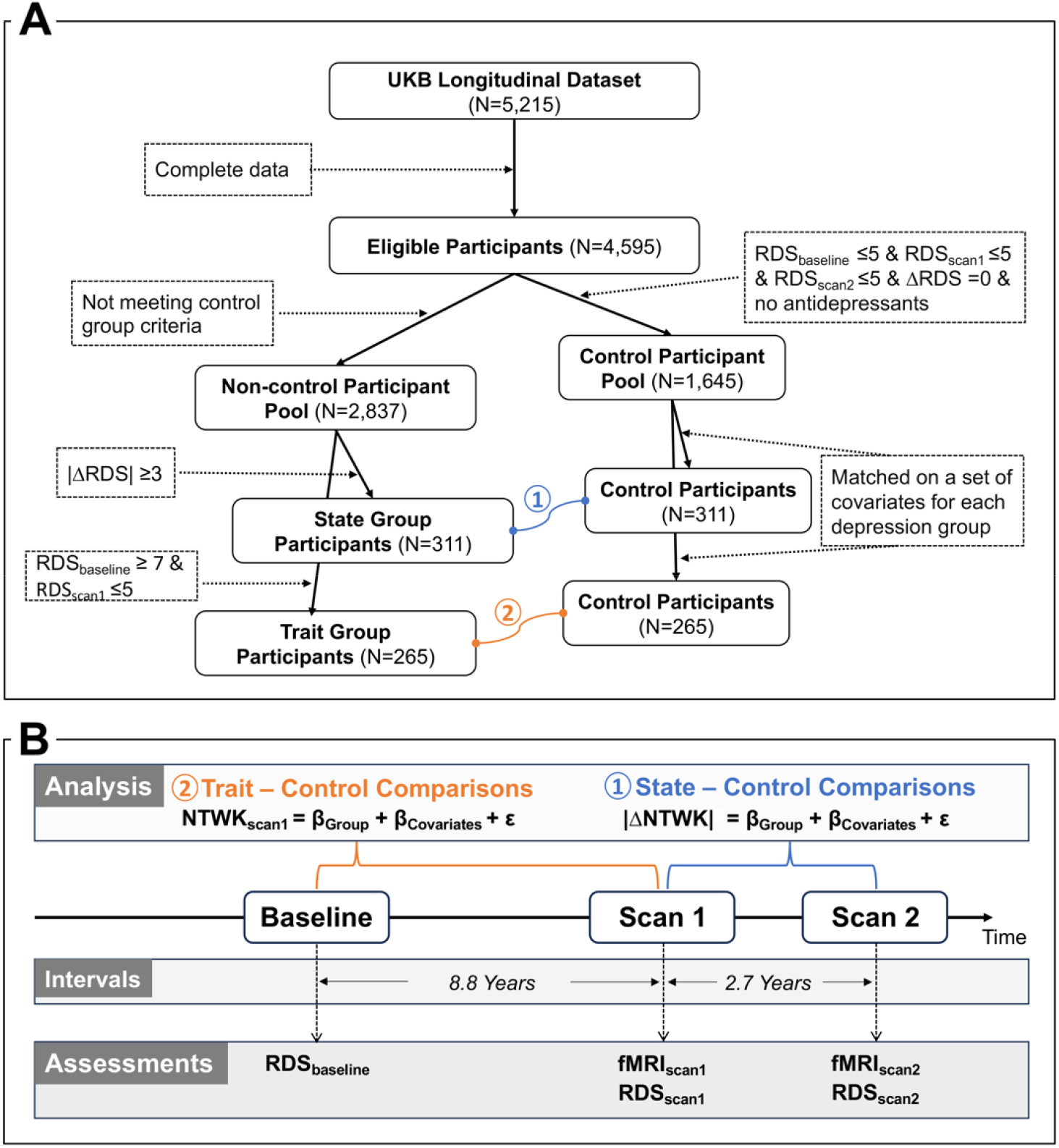
Flowchart of the study sample (A) and schema of analysis pipeline (B). Specifically, the RDS_baseline_, RDS_scan1_, and RDS_scan2_ represent sum scores of Recent Depressive Symptoms (RDS) obtained at different time points, with subscripts indicating the assessment time, whereas |ΔRDS| denotes the absolute longitudinal change score of RDS between two neuroimaging scans. The final sample consisted of two pairs of matched groups, connected by curved lines. Group comparisons between the matched state-control (⍰) and trait-control (⍰) were performed separately, utilizing brain network measures (NTWK) assessed at different time points. Importantly, in trait-control comparisons, the network measures at scan1 (NTWK_scan1_) were considered dependent variables, while in state-control comparisons, the absolute values of longitudinal changes in the network measures (|ΔNTWK|), were included as dependent variables. All these dependent variables were modeled as a function of the group variable (e.g., state vs. control), while accounting for covariates.

### 2. Defining Depression and Control Groups

#### 2.1. State depression group

Longitudinal data were used to capture changes in state depression, identifying participants with a large (≥3 points) absolute change in longitudinal score of Recent Depressive Symptoms (RDS) between the first and follow-up imaging assessments (i.e., scan1 and scan2). The absolute change (|ΔRDS|) was chosen to allow for symptom severity changes in both increased and decreased directions, capturing the full spectrum of state-dependent fluctuations. The RDS is a 4-item scale with scores ranging from 4 to 16, validated against standardized scales of depression (Dutt et al., 2022). The 3-point change threshold, representing 25% of the full 12-point scale for RDS, signifies clinically meaningful changes in state depression while maximizing sample size (see Supplemental Materials ***Table S1***). As a reference, a 25% change on the PHQ scale (0-27) would exceed the 5-point cut-off for mild depressive symptoms (Kroenke et al., 2001). N=311 participants met the |ΔRDS| >3 inclusion requirement for the state depression group.

### 2.2 Trait depression group

Trait depression, reflecting a long-term tendency to experiencing depressive symptoms (Klein et al., 2011) was operationalized in our study as individuals with prior depression but in remission at the time of scanning (i.e., scan1). Specifically, participants with an RDS score at the baseline equal to or above 7 (RDS_baseline_ ≥ 7) AND an RDS score at the first imaging assessment equal to or below 5 (RDS_scan1_ ≤ 5) were included in the trait depression group. Notably, our previous work has mapped RDS onto PHQ-9 (Dutt et al., 2022), where an RDS score of 7 corresponds to PHQ = 5, the clinical threshold for mild depression (Kroenke et al., 2001). Additionally, an RDS score of 5 represents a minimal degree of depression and RDS_scan1_ ≤ 5 indicates a remission status at the first imaging assessment. N=265 participants met the RDS_baseline_ ≥ 7 & RDS_scan1_ ≤ 5 inclusion requirements for the trait depression group.

### 2.3. Control participants and group matching

A pool of potential control participants was identified, meeting criteria of minimal depression scores across all three time points (RDS_baseline_ ≤ 5 & RDS_scan1_ ≤ 5 & RDS_scan2_ ≤ 5), and no change in depression score between scan 1 and scan 2 (ΔRDS = 0). These individuals were not on antidepressant medication at any time point either, ensuring a healthy reference group (see list for antidepressants in Supplemental Materials). We identified a total of N=1,448 participants meeting these criteria. From this total pool, we separately selected the equivalent number of participants for both the state and trait depression groups. These selections were made to ensure matching for each depression group on sex, age, in-scanner head motion (i.e., averaged relative framewise displacement), scanning site, and alcohol intake frequency at the first imaging assessment (i.e., at scan1), thus minimizing potential confounding effects on between-group differences under investigation.

Matches were identified using propensity scores with an optimal matching algorithm, selected from several algorithms (see matching diagnostics in Supplemental Materials Table S3). The matched groups demonstrated improved covariate balance for all variables of interest, with no significant group difference observed between the depression groups and their respective matched controls (all p’s>0.05; Supplemental Materials **Table S4**).

### 3. fMRI data acquisition and preprocessing

The UKB acquired resting-state neuroimaging data using a 3T Siemens Skyra (2.4mm isotropic voxel size, TR=0.735 sec, multiband factor 8). The detailed scanning protocols are documented online (https://biobank.ctsu.ox.ac.uk/crystal/refer.cgi?id=2367). Our study used preprocessed data that were released via the UKB showcase. The preprocessing steps included distortion correction, motion correction, high pass temporal filtering, and BOLD signal denoising using ICA-FIX (details in (Alfaro-Almagro et al., 2018).

#### 3.1. Brain networks identified by probabilistic functional modes estimation

Probabilistic Functional Modes (PROFUMO) is a hierarchical Bayesian approach to decompose resting state neuroimaging data into a set of modes, representing resting-state brain networks (Harrison et al., 2015, 2020) . Each mode is described by a spatial map, network matrix (i.e., connectivity matrix - estimated as the partial correlation between mode time-courses), and mode amplitude. Each of these PROFUMO outputs is estimated separately per longitudinal scan and simultaneously at the group and individual levels. PROFUMO offers advantages in effectively capturing spatial overlaps in network structures that are indicative of shared spatial organization across networks (Bijsterbosch et al., 2019), and demonstrates increased sensitivity in discerning individual-specific network configuration (Bijsterbosch et al., 2018; Harrison et al., 2020). To maintain consistency across state and trait analyses, we merged resting-state fMRI data from all four groups, resulting in a sample size of 1,030 unique individuals. Each participant contributed data from two time points (scan1 and scan2), totaling 2,060 scans for the final PROFUMO decomposition.

In the main analysis, we set the PROFUMO mode dimension to 15, aligning with our focus on large-scale networks. After removing two spurious modes (Supplemental Materials), we included 13 meaningful brain networks for statistical analyses and used each of the three PROFUMO outputs per network as dependent variables. To ensure robustness, we also examined 10 and 20 dimensions, validating significant findings from the 15-dimensional outputs. This adjustment from our preregistration, originally focused on 20-dimensional networks, aimed to streamline multiple-testing comparisons, and emphasize a low-dimensional decomposition into large-scale canonical resting-state networks.

### 4. Statistical Analysis

Using linear regression models, separate analyses compared group differences in neural correlates between each of the two depression groups and their respective control groups. Specifically, state-control comparisons assessed longitudinal changes in network measures between two neuroimaging scans, while trait-control comparisons considered network measures at scan1 as the dependent variables (see ***Figure 1B***). These dependent variables comprised PROFUMO outputs, including mode amplitude (indicating overall signal fluctuation), network matrix (reflecting partial correlation between mode timeseries), and spatial overlap matrix (illustrating full correlation between mode spatial maps).

#### 4.1 Main Analyses

The primary analysis for state depression involved computing longitudinal changes in each of the three brain network measures (i.e., PROFUMO outputs) between two scans (i.e., scan2 minus scan1). The absolute values of these changes served as dependent variables in subsequent group-comparison analyses. In the main analysis for trait depression, the network measures at scan1 were utilized as dependent variables in the group-comparison analyses. Thus, separate regression analyses were conducted for state-control and trait-control group comparisons, utilizing different dependent variables (see analysis equations in ***Figure 1B***). Within each comparison, we performed statistical analysis separately for each class of PROFUMO outputs. Covariates, including sex, age, in-scanner head motion, scanning site, alcohol intake frequency, time interval between two scans (only for state-control comparisons), and use of antidepressants, were adjusted in all analyses.

To address multiple testing concerns within each class of three PROFUMO outputs, False Discovery Rate (FDR) correction was applied. This approach was chosen to accommodate distinct hypotheses for different PROFUMO outputs.

The number of separate univariate analyses was determined by the PROFUMO dimensionality. In our main analyses (using 13 meaningful modes), we obtained 13 mode amplitudes, 78 temporal edges from connectivity matrix, and 78 spatial edges indicative of spatial overlaps between modes from spatial correlation matrix. These edges represent unique pairwise correlation coefficients derived from the upper or low triangle of the connectivity or spatial correlation matrices (i.e., 13*12/2=78), constructed from the temporal or spatial correlations between each pair of timeseries or spatial maps of nodes.

#### 4.2. Follow-up Analyses

Several follow-up analyses were conducted to further validate the statistical significance of the group comparison results from our main analyses.

First, we performed regression analyses to investigate symptom magnitude-dependent effects within depression groups in case significant group differences were observed in any of the three PROFUMO outputs. This involved separate analyses for each significant result from the group comparisons among individuals within the relevant depression group. For example, if individuals with trait depression exhibited larger connectivity strength than their control counterparts, the subsequent tests would examine whether, among individuals with trait depression, greater connectivity strength was associated with higher symptom severity.

Secondly, we conducted follow-up analyses aiming to identify robust findings across different dimensionalities. We first identified the ‘best-matched’ mode(s) from 10- and 20-dimension decompositions for the target modes from the 15-dimension decomposition, using Cosine similarity coefficients (see details in Supplemental Materials). Subsequently, we repeated group comparisons on the identified modes from 10 and 20 dimensionalities to confirm the persistence of significant results observed in the 15-dimensionality. While our pre-registration suggested the Hungarian algorithm (Kuhn, 1955) for mode matching, we adjusted our strategy to prioritize replicating findings for specific modes, favoring spatial similarity over the complete data matching. This choice prevents issues with imbalanced dimensionalities (e.g., 10 vs. 15), ensuring effective mode matching.

## Results

### 1. Characteristics of Participants in Each Group

With our operationalized definitions of the state and trait depression groups, participants in the state group showed higher RDS scores across all time points, whereas participants in the trait group only reported higher RDS scores compared to controls at baseline and scan 2 (p’s < 0.0001), with comparable RDS scores to controls at scan 1, indicating the intended remission status (***Table 1***).

Importantly, these operational definitions achieved the desired state-trait dissociation by reducing the known high correlations between these two constructs. Specifically, for the participants in the state group, the longitudinal changes in RDS between two scans (ΔRDS), reflecting state-dependent depression experiences, showed a negligible correlation (r=-0.01) with the baseline RDS (RDS_baseline_). This substantially reduced the initial correlation between the ΔRDS and RDS_baseline_ (r=0.291) in the full UKB sample (n=4,595). A similar state-trait dissociation was observed for the participants in the trait depression group: The RDS_baseline_, used to define trait experiences, exhibited only a correlation of r=0.03 with the RDS at scan1 (RDS_scan1_), compared to the initial r=0.52 in the full UKB sample. The RDS_scan1_ here indexed the present depression symptoms of the participants from the trait depression group at the time of scan1, when the brain network measures were assessed for these participants.

Furthermore, the control participants exhibited similar time intervals between different assessments, along with demographic and several other variables that were matched between the depression and control groups (**Table 1**; Supplemental Materials **Table S4**).

#### 2. Decomposed PROFUMO Modes

At the group level, a total of 15 probabilistic modes were estimated across all depression and control participants using both scan1 and scan2 data, 13 of which were identified as large-scale brain networks representing meaningful brain signals (see full decomposition in Supplemental Materials ***Figure S1***). The spatial distributions of these probabilistic modes highly resembled canonical brain networks including the default mode, frontoparietal, visual and motor networks. Similar resemblances were also observed in 10- and 20-dimension decompositions, except that some of the networks in one decomposition appeared to merge into one or split into two or more modes in another decomposition (Supplemental Materials ***Figure S2-3***).

### 3. Group Comparisons in Neural Correlates

Individuals experiencing trait depression exhibited significantly higher amplitude in two visual networks in contrast to the matched control participants (***Figure 2***). This effect was robust against the inclusion of all covariates including antidepressant usage (β_visual1_=0.045, β_visual2_=0.034, FDR corrected p’s<0.05). These findings also replicated in the follow-up analyses using PROFUMO outputs from the 10- and 20-dimension decompositions (β’s > 0.02, p’s < 0.03; Supplemental Results ***Table S5***). However, the magnitude of the mode amplitude within the visual networks was not associated with depression symptom severity at baseline among individuals within the trait group (β=-0.009, p>0.8). Further, we did not find significant group differences in spatial overlaps or network matrices for the trait analyses (all FDR corrected p’s>0.05).

**Figure 2.**
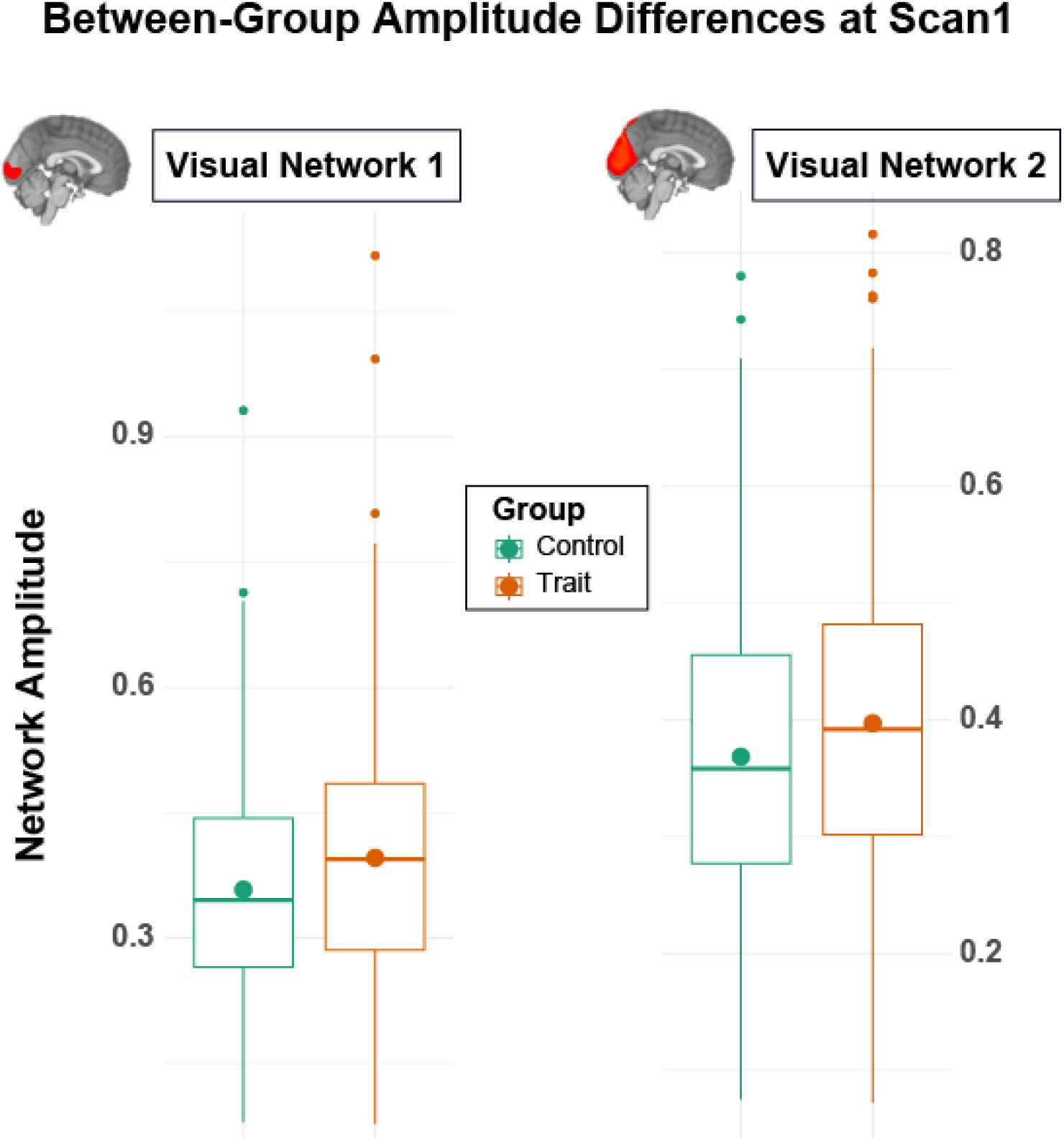
Box plots showing group differences in amplitude of two visual networks at the time of scan1, between individuals with trait experience of depression (in orange) and control participants (in green), with higher mean amplitudes in both networks, as annotated by filled circles, in the trait depression group. Note, separate Y-axis scales were used to highlight the group mean differences within each comparison.

After corrections for multiple testing and potential confounding effects, we did not find significant group differences in any absolute longitudinal changes in PROFUMO outputs (amplitude, network matrix and spatial overlap) between individuals from the state-depression group and the matched controls (all FDR corrected p’s>0.05).

## Discussion

In the current study, we designed group contrast that effectively disassociated the neural correlates of state and trait experiences of depression. Using matched control participants as the reference, our results showed that greater amplitudes in two visual networks were associated with trait depression and that these associative effects were robust against relevant confounders and across different dimensionalities of brain networks. These findings demonstrate that the overall BOLD signal fluctuations within the visual networks may serve as a potential biomarker for trait depression.

### Visual cortex and trait depression

Depression is a mental disorder characterized by dysfunctional brain networks (Williams, 2016). Although the majority of the Major Depressive Disorder (MDD) literature focuses on alterations in the default mode, salience, and central executive networks (Kaiser et al., 2015), there is increasing evidence linking depression to structural and functional alterations in sensory and motor networks, including visual networks (Wu et al., 2023; Zhukovsky et al., 2021).

Structurally, individuals with MDD often exhibit increased volume (Ancelin et al., 2019), greater asymmetry (Maller et al., 2014), and altered surface area (Schmaal et al., 2017) in the occipital cortex compared to healthy controls. These structural alterations have been further associated with symptom recurrence and duration in MDD patients (Y. Kang et al., 2023). Functionally, MDD patients demonstrate various abnormalities within the visual network. This includes decreased regional homogeneity (Peng et al., 2011; M. Yan et al., 2021), lower nodal efficiency in occipital areas (Xu et al., 2022), aberrant resting-state functional connectivity within the network (Lu et al., 2020), and disruptions in connections between the visual network and other brain networks (Kaiser et al., 2015; C.-G. Yan et al., 2019; Young et al., 2023). Moreover, depressed individuals exhibit abnormal filtering of irrelevant information in the visual cortex (Desseilles et al., 2009) and altered activity patterns in visual association areas affecting working memory (Le et al., 2017). These findings collectively underscore the crucial role of the visual cortex in the altered visual perception commonly observed in MDD patients (Friberg & Borrero, 2000; Salmela et al., 2021). Such alterations may be attributed to reduced concentrations of the neurotransmitter GABA in MDD patients, leading to deficits in the inhibition or suppression of relevant visual information processing (Price et al., 2009; Song et al., 2021). Our findings are furthermore consistent with findings of increased amplitude of low-frequency fluctuation (ALFF) in the occipital cortex in patients with remitted but not current depression, in contrast to healthy controls (Cheng et al., 2019).

Yet, our study found no association between the magnitude of amplitude increase and symptom severity among individuals with trait depression, likely due to limited variability in symptom severity within this group. Nearly half of the individuals (120 out of 265) in this depression group exhibited mild-level symptoms (RDS = 7), which was the minimum criterion for inclusion in the trait group. However, despite the limited variability, the propensity for depression was sufficient at the group level to be associated with differences in brain function, evident in the between-group differences in visual network amplitude.

### Null findings for state depression

In contrast to our hypothesis, no significant differences in network connectivity or other measures were found between the state depression group and the control group. These null findings may have resulted from several factors.

Firstly, the state experience in our study was defined as substantial fluctuations in symptom severity between two imaging scans. While the overall symptom severity score provides insight into the general experiences of state depression, it may not capture fluctuations in specific symptoms. This limitation makes the investigation into associated brain measures susceptible to individual heterogeneity, as network measures for different symptoms at the individual level may not converge at the group level.

Secondly, we opted for a cut-off score of 3 to define the state experience, aiming at capturing meaningful changes in symptoms, equivalent to 25% of the full 12-point scale. This choice was made to also maximize the sample size with sufficient statistical power to ensure robustness in detecting effects of interest. For example, increasing the cut-off score to 4, would have reduced the combined sample size to 256 (i.e., n=311 to n=128 for trait depression and control groups each; Supplemental Materials ***Table S1***), potentially leading to underpowered analyses. Retrospective power analysis using the reduced sample size showed 70% power to detect a small group effect at β=0.05, slightly higher than β_visual1_=0.045 in our main findings, compared to the 93% power obtained from the current combined n=622. However, it is possible that this mild cut-off may have been insufficient to detect state-related alterations in network connectivity or other brain measures.

Additionally, our current definition of state depression considered the magnitude of symptom fluctuations, combining both positive and negative changes in symptom severity between two imaging scans. If different brain networks are engaged in different directions of state changes, our approach might fail to detect group-level effects due to a potential mixture of results. Unfortunately, our sample size for the state depression participants was nearly halved when considering two separate groups with opposing change directions (n=154 for negative changes and n=157 for positive changes), leaving separate testing underpowered.

Lastly, given the dynamic nature of state depression experiences, the use of evoked study designs might offer greater sensitivity in capturing alterations in state-related neural correlates. Recent meta-analysis studies seem to support this notion, revealing reduced brain activation in a wide range of cortical and subcortical regions for MDD patients during emotional processing after antidepressant treatment in contrast to the baseline (Delaveau et al., 2011). Additionally, altered activity patterns have been observed in tasks related to emotional processing or executive functioning tasks, or across aggregated tasks from these two domains (Gray et al., 2020).

### Null findings in the default mode network (DMN)

Our current investigation did not find DMN differences in any PROFUMO output for either the trait or state comparison. This seemingly surprising result contrasts with the prior implications of this brain network in relation to depression. It is important to note, however, that previous research has also yielded inconsistent and/or contradictory findings regarding the role of the DMN in MDD (Javaheripour et al., 2021; Kaiser et al., 2015; Mulders et al., 2015; C.-G. Yan et al., 2019). This inconsistency underscores the challenges in identifying robust neural correlates of depression across studies.

Despite employing a data-driven approach to identify brain networks including the DMN (i.e., mode 5 in **Figure S1**), and utilizing data from a large-scale prospective epidemiological resource (Sudlow et al., 2015) to dissociate neural correlates of state and trait depression, our null findings align with the broader inconsistencies observed in the literature regarding alterations in the DMN and other brain networks in previous studies.

### Limitations

The current study had several limitations. Firstly, our study sample is a subset from the UK Biobank study, comprising specific age groups (i.e., middle to older adults) and predominantly White (Sudlow et al., 2015). This limits the generalization of our findings to more diverse populations with different demographic characteristics. Secondly, while we carefully generated two control groups and matched them for each of the target depression groups based on a set of crucial confounding variables, our analyses for group comparisons did not include psychosocial or lifestyle factors that can contribute to individual variations in depression symptoms (Aguilar-Latorre et al., 2022; Remes et al., 2021; Sarris et al., 2014; Zhao et al., 2023). These additional factors may also influence the associative effects under investigation, and future studies should consider integrating them into analysis. Lastly, although we employed a data-driven approach to identify large-scale brain networks, this method appeared to favor the discovery of networks involving cortical regions over subcortical ones. These networks showed higher signal intensities predominantly in cortical areas, likely due to higher signal-to-noise ratios. Given the documented structural (Ho et al., 2020; Schmaal et al., 2016) and functional alterations (Gray et al., 2020; Miller et al., 2015; Xiong et al., 2021) in subcortical regions in MDD, it may be prudent to explore supplementary approaches to better detect brain networks involving subcortical regions in the future.

## Conclusion

Incorporating pre-registered hypotheses and methods, this study aimed to examine the potentially distinct neural correlates of state and trait experiences of depression. We observed significantly higher amplitude in two visual networks for individuals in the trait depression group compared to controls. No significant differences in network measures were found in relation to state depression . Our findings suggest potentially altered visual information processing for individuals with a persistent tendency to experience depressive symptoms. These results highlight the potential contribution of visual networks to the psychopathology of depression that has been largely overlooked in the literature and provide evidence for neural correlates specific to trait experiences of depression.

## Reference

1. Admon, R., Holsen, L. M., Aizley, H., Remington, A., Whitfield-Gabrieli, S., Goldstein, J. M., & Pizzagalli, D. A. (2015). Striatal Hypersensitivity During Stress in Remitted Individuals with Recurrent Depression. Biological Psychiatry, 78(1), 67–76. 10.1016/j.biopsych.2014.09.019

2. Aguilar-Latorre, A., Algorta, G. P., Navarro-Guzmán, C., Serrano-Ripoll, M. J., & Oliván-Blázquez, B. (2022). Effectiveness of a lifestyle modification programme in the treatment of depression symptoms in primary care. Frontiers in Medicine, 9. 10.3389/fmed.2022.954644

3. Alfaro-Almagro, F., Jenkinson, M., Bangerter, N. K., Andersson, J. L. R., Griffanti, L., Douaud, G., Sotiropoulos, S. N., Jbabdi, S., Hernandez-Fernandez, M., Vallee, E., Vidaurre, D., Webster, M., McCarthy, P., Rorden, C., Daducci, A., Alexander, D. C., Zhang, H., Dragonu, I., Matthews, P. M., … Smith, S. M. (2018). Image processing and Quality Control for the first 10,000 brain imaging datasets from UK Biobank. NeuroImage, 166, 400–424. 10.1016/j.neuroimage.2017.10.034

4. Ancelin, M.-L., Carrière, I., Artero, S., Maller, J., Meslin, C., Ritchie, K., Ryan, J., & Chaudieu, I. (2019). Lifetime major depression and grey-matter volume. Journal of Psychiatry & NeurosciencelJ: JPN, 44(1), 45–53. 10.1503/jpn.180026

5. Bijsterbosch, J. D., Beckmann, C. F., Woolrich, M. W., Smith, S. M., & Harrison, S. J. (2019). The relationship between spatial configuration and functional connectivity of brain regions revisited. eLife, 8, e44890. 10.7554/eLife.44890

6. Bijsterbosch, J. D., Woolrich, M. W., Glasser, M. F., Robinson, E. C., Beckmann, C. F., Van Essen, D. C., Harrison, S. J., & Smith, S. M. (2018). The relationship between spatial configuration and functional connectivity of brain regions. eLife, 7, e32992. 10.7554/eLife.32992

7. Braund, T. A., Breukelaar, I. A., Griffiths, K., Tillman, G., Palmer, D. M., Bryant, R., Phillips, M. L., Harris, A. W. F., & Korgaonkar, M. S. (2022). Intrinsic Functional Connectomes Characterize Neuroticism in Major Depressive Disorder and Predict Antidepressant Treatment Outcomes. Biological Psychiatry. Cognitive Neuroscience and Neuroimaging, 7(3), 276–284. 10.1016/j.bpsc.2021.07.010

8. Cheng, C., Dong, D., Jiang, Y., Ming, Q., Zhong, X., Sun, X., Xiong, G., Gao, Y., & Yao, S. (2019). State-Related Alterations of Spontaneous Neural Activity in Current and Remitted Depression Revealed by Resting-State fMRI. Frontiers in Psychology, 10, 245. 10.3389/fpsyg.2019.00245

9. Delaveau, P., Jabourian, M., Lemogne, C., Guionnet, S., Bergouignan, L., & Fossati, P. (2011). Brain effects of antidepressants in major depression: A meta-analysis of emotional processing studies. Journal of Affective Disorders, 130(1–2), 66–74. 10.1016/j.jad.2010.09.032

10. Desseilles, M., Balteau, E., Sterpenich, V., Dang-Vu, T. T., Darsaud, A., Vandewalle, G., Albouy, G., Salmon, E., Peters, F., Schmidt, C., Schabus, M., Gais, S., Degueldre, C., Phillips, C., Luxen, A., Ansseau, M., Maquet, P., & Schwartz, S. (2009). Abnormal neural filtering of irrelevant visual information in depression. The Journal of Neuroscience: The Official Journal of the Society for Neuroscience, 29(5), 1395–1403. 10.1523/JNEUROSCI.3341-08.2009

11. Dinga, R., Schmaal, L., Penninx, B. W. J. H., van Tol, M. J., Veltman, D. J., van Velzen, L., Mennes, M., van der Wee, N. J. A., & Marquand, A. F. (2019). Evaluating the evidence for biotypes of depression: Methodological replication and extension of. NeuroImage. Clinical, 22, 101796. 10.1016/j.nicl.2019.101796

12. Drysdale, A. T., Grosenick, L., Downar, J., Dunlop, K., Mansouri, F., Meng, Y., Fetcho, R. N., Zebley, B., Oathes, D. J., Etkin, A., Schatzberg, A. F., Sudheimer, K., Keller, J., Mayberg, H. S., Gunning, F. M., Alexopoulos, G. S., Fox, M. D., Pascual-Leone, A., Voss, H. U., … Liston, C. (2017). Resting-state connectivity biomarkers define neurophysiological subtypes of depression. Nature Medicine, 23(1), 28–38. 10.1038/nm.4246

13. Dutt, R. K., Hannon, K., Easley, T. O., Griffis, J. C., Zhang, W., & Bijsterbosch, J. D. (2022). Mental health in the UK Biobank: A roadmap to self-report measures and neuroimaging correlates. Human Brain Mapping, 43(2), 816–832. 10.1002/hbm.25690

14. Flint, C., Cearns, M., Opel, N., Redlich, R., Mehler, D. M. A., Emden, D., Winter, N. R., Leenings, R., Eickhoff, S. B., Kircher, T., Krug, A., Nenadic, I., Arolt, V., Clark, S., Baune, B. T., Jiang, X., Dannlowski, U., & Hahn, T. (2021). Systematic misestimation of machine learning performance in neuroimaging studies of depression. Neuropsychopharmacology, 46(8), Article 8. 10.1038/s41386-021-01020-7

15. Fournier, J. C., Chase, H. W., Greenberg, T., Etkin, A., Almeida, J. R., Stiffler, R., Deckersbach, T., Weyandt, S., Cooper, C., Toups, M., Carmody, T., Kurian, B., Peltier, S., Adams, P., McInnis, M. G., Oquendo, M. A., McGrath, P. J., Fava, M., Weissman, M., … Phillips, M. L. (2017). Neuroticism and Individual Differences in Neural Function in Unmedicated Major Depression: Findings from the EMBARC Study. Biological Psychiatry. Cognitive Neuroscience and Neuroimaging, 2(2), 138–148. 10.1016/j.bpsc.2016.11.008

16. Friberg, T. R., & Borrero, G. (2000). Diminished perception of ambient light: A symptom of clinical depression? Journal of Affective Disorders, 61(1–2), 113–118. 10.1016/s0165-0327(99)00194-9

17. Friedrich, M. J. (2017). Depression Is the Leading Cause of Disability Around the World. JAMA, 317(15), 1517. 10.1001/jama.2017.3826

18. Gray, J. P., Müller, V. I., Eickhoff, S. B., & Fox, P. T. (2020). Multimodal Abnormalities of Brain Structure and Function in Major Depressive Disorder: A Meta-Analysis of Neuroimaging Studies. American Journal of Psychiatry, 177(5), 422–434. 10.1176/appi.ajp.2019.19050560

19. Greene, A. S., Shen, X., Noble, S., Horien, C., Hahn, C. A., Arora, J., Tokoglu, F., Spann, M. N., Carrión, C. I., Barron, D. S., Sanacora, G., Srihari, V. H., Woods, S. W., Scheinost, D., & Constable, R. T. (2022). Brain–phenotype models fail for individuals who defy sample stereotypes. Nature, 609(7925), Article 7925. 10.1038/s41586-022-05118-w

20. Group, F.-N. B. W. (2021). FDA-NIH Biomarker Working Group. In BEST (Biomarkers, EndpointS, and other Tools) Resource [Internet]. Food and Drug Administration (US). https://www.ncbi.nlm.nih.gov/books/NBK338449/

21. Hannon, K., Easley, T., Zhang, W., Lew, D., Thornton, V., Sotiras, A., Sheline, Y. I., Marquand, A., Barch, D. M., & Bijsterbosch, J. D. (2022). Heterogeneity in Depression: Evidence for distinct clinical and neurobiological profiles (p. 2022.12.07.22283225). medRxiv. 10.1101/2022.12.07.22283225

22. Harrison, S. J., Bijsterbosch, J. D., Segerdahl, A. R., Fitzgibbon, S. P., Farahibozorg, S.-R., Duff, E. P., Smith, S. M., & Woolrich, M. W. (2020). Modelling subject variability in the spatial and temporal characteristics of functional modes. NeuroImage, 222, 117226. 10.1016/j.neuroimage.2020.117226

23. Harrison, S. J., Woolrich, M. W., Robinson, E. C., Glasser, M. F., Beckmann, C. F., Jenkinson, M., & Smith, S. M. (2015). Large-scale Probabilistic Functional Modes from resting state fMRI. NeuroImage, 109, 217–231. 10.1016/j.neuroimage.2015.01.013

24. Ho, T. C., Gutman, B., Pozzi, E., Grabe, H. J., Hosten, N., Wittfeld, K., Völzke, H., Baune, B., Dannlowski, U., Förster, K., Grotegerd, D., Redlich, R., Jansen, A., Kircher, T., Krug, A., Meinert, S., Nenadic, I., Opel, N., Dinga, R., … Schmaal, L. (2020). Subcortical shape alterations in major depressive disorder: Findings from the ENIGMA major depressive disorder working group. Human Brain Mapping, 43(1), 341–351. 10.1002/hbm.24988

25. Javaheripour, N., Li, M., Chand, T., Krug, A., Kircher, T., Dannlowski, U., Nenadić, I., Hamilton, J. P., Sacchet, M. D., Gotlib, I. H., Walter, H., Frodl, T., Grimm, S., Harrison, B. J., Wolf, C. R., Olbrich, S., van Wingen, G., Pezawas, L., Parker, G., … Wagner, G. (2021). Altered resting-state functional connectome in major depressive disorder: A mega-analysis from the PsyMRI consortium. Translational Psychiatry, 11(1), Article 1. 10.1038/s41398-021-01619-w

26. Kaiser, R. H., Andrews-Hanna, J. R., Wager, T. D., & Pizzagalli, D. A. (2015). Large-Scale Network Dysfunction in Major Depressive Disorder: A Meta-analysis of Resting-State Functional Connectivity. JAMA Psychiatry, 72(6), 603–611. 10.1001/jamapsychiatry.2015.0071

27. Kang, L., Zhang, A., Sun, N., Liu, P., Yang, C., Li, G., Liu, Z., Wang, Y., & Zhang, K. (2018). Functional connectivity between the thalamus and the primary somatosensory cortex in major depressive disorder: A resting-state fMRI study. BMC Psychiatry, 18(1), 339. 10.1186/s12888-018-1913-6

28. Kang, Y., Kang, W., Kim, A., Tae, W.-S., Ham, B.-J., & Han, K.-M. (2023). Decreased cortical gyrification in major depressive disorder. Psychological Medicine, 53(16), 7512–7524. 10.1017/S0033291723001216

29. Kennis, M., Gerritsen, L., van Dalen, M., Williams, A., Cuijpers, P., & Bockting, C. (2020). Prospective biomarkers of major depressive disorder: A systematic review and meta-analysis. Molecular Psychiatry, 25(2), 321–338. 10.1038/s41380-019-0585-z

30. Klein, D. N., Kotov, R., & Bufferd, S. J. (2011). Personality and Depression: Explanatory Models and Review of the Evidence. Annual Review of Clinical Psychology, 7, 269–295. 10.1146/annurev-clinpsy-032210-104540

31. Kroenke, K., Spitzer, R. L., & Williams, J. B. (2001). The PHQ-9: Validity of a brief depression severity measure. Journal of General Internal Medicine, 16(9), 606–613. 10.1046/j.1525-1497.2001.016009606.x

32. Kuhn, H. W. (1955). The Hungarian method for the assignment problem. Naval Research Logistics Quarterly, 2(1–2), 83–97. 10.1002/nav.3800020109

33. Le, T. M., Borghi, J. A., Kujawa, A. J., Klein, D. N., & Leung, H.-C. (2017). Alterations in visual cortical activation and connectivity with prefrontal cortex during working memory updating in major depressive disorder. NeuroImagelJ: Clinical, 14, 43–53. 10.1016/j.nicl.2017.01.004

34. Lu, F., Cui, Q., Huang, X., Li, L., Duan, X., Chen, H., Pang, Y., He, Z., Sheng, W., Han, S., Chen, Y., Yang, Y., Luo, W., Yu, Y., Jia, X., Tang, Q., Li, D., Xie, A., & Chen, H. (2020). Anomalous intrinsic connectivity within and between visual and auditory networks in major depressive disorder. Progress in Neuro-Psychopharmacology and Biological Psychiatry, 100, 109889. 10.1016/j.pnpbp.2020.109889

35. Maller, J. J., Thomson, R. H. S., Rosenfeld, J. V., Anderson, R., Daskalakis, Z. J., & Fitzgerald, P. B. (2014). Occipital bending in depression. Brain, 137(6), 1830–1837. 10.1093/brain/awu072

36. Miller, C. H., Hamilton, J. P., Sacchet, M. D., & Gotlib, I. H. (2015). Meta-analysis of Functional Neuroimaging of Major Depressive Disorder in Youth. JAMA Psychiatry, 72(10), 1045–1053. 10.1001/jamapsychiatry.2015.1376

37. Ming, Q., Zhong, X., Zhang, X., Pu, W., Dong, D., Jiang, Y., Gao, Y., Wang, X., Detre, J. A., Yao, S., & Rao, H. (2017). State-Independent and Dependent Neural Responses to Psychosocial Stress in Current and Remitted Depression. The American Journal of Psychiatry, 174(10), 971–979. 10.1176/appi.ajp.2017.16080974

38. Mulders, P. C., van Eijndhoven, P. F., Schene, A. H., Beckmann, C. F., & Tendolkar, I. (2015). Resting-state functional connectivity in major depressive disorder: A review. Neuroscience and Biobehavioral Reviews, 56, 330–344. 10.1016/j.neubiorev.2015.07.014

39. Oathes, D. J., Patenaude, B., Schatzberg, A. F., & Etkin, A. (2015). Neurobiological signatures of anxiety and depression in resting-state functional magnetic resonance imaging. Biological Psychiatry, 77(4), 385–393. 10.1016/j.biopsych.2014.08.006

40. Peng, D.-H., Jiang, K., Fang, Y.-R., Xu, Y.-F., Shen, T., Long, X.-Y., Liu, J., & Zang, Y.-F. (2011). Decreased regional homogeneity in major depression as revealed by resting-state functional magnetic resonance imaging. Chinese Medical Journal, 124(3), 369–373.

41. Price, R. B., Shungu, D. C., Mao, X., Nestadt, P., Kelly, C., Collins, K. A., Murrough, J. W., Charney, D. S., & Mathew, S. J. (2009). Amino Acid Neurotransmitters Assessed by Proton Magnetic Resonance Spectroscopy: Relationship to Treatment Resistance in Major Depressive Disorder. Biological Psychiatry, 65(9), 792–800. 10.1016/j.biopsych.2008.10.025

42. Ray, D., Bezmaternykh, D., Mel’nikov, M., Friston, K. J., & Das, M. (2021). Altered effective connectivity in sensorimotor cortices is a signature of severity and clinical course in depression. Proceedings of the National Academy of Sciences, 118(40), e2105730118. 10.1073/pnas.2105730118

43. Remes, O., Mendes, J. F., & Templeton, P. (2021). Biological, Psychological, and Social Determinants of Depression: A Review of Recent Literature. Brain Sciences, 11(12), 1633. 10.3390/brainsci11121633

44. Saberi, A., Mohammadi, E., Zarei, M., Eickhoff, S. B., & Tahmasian, M. (2022). Structural and functional neuroimaging of late-life depression: A coordinate-based meta-analysis. Brain Imaging and Behavior, 16(1), 518–531. 10.1007/s11682-021-00494-9

45. Salmela, V., Socada, L., Söderholm, J., Heikkilä, R., Lahti, J., Ekelund, J., & Isometsä, E. (2021). Reduced visual contrast suppression during major depressive episodes. Journal of Psychiatry & NeurosciencelJ: JPN, 46(2), E222–E231. 10.1503/jpn.200091

46. Sarris, J., O’Neil, A., Coulson, C. E., Schweitzer, I., & Berk, M. (2014). Lifestyle medicine for depression. BMC Psychiatry, 14, 107. 10.1186/1471-244X-14-107

47. Schmaal, L. (2022). The Search for Clinically Useful Neuroimaging Markers of Depression—A Worthwhile Pursuit or a Futile Quest? JAMA Psychiatry, 79(9), 845–846. 10.1001/jamapsychiatry.2022.1606

48. Schmaal, L., Hibar, D. P., Sämann, P. G., Hall, G. B., Baune, B. T., Jahanshad, N., Cheung, J. W., van Erp, T. G. M., Bos, D., Ikram, M. A., Vernooij, M. W., Niessen, W. J., Tiemeier, H., Hofman, A., Wittfeld, K., Grabe, H. J., Janowitz, D., Bülow, R., Selonke, M., … Veltman, D. J. (2017). Cortical abnormalities in adults and adolescents with major depression based on brain scans from 20 cohorts worldwide in the ENIGMA Major Depressive Disorder Working Group. Molecular Psychiatry, 22(6), 900–909. 10.1038/mp.2016.60

49. Schmaal, L., Veltman, D. J., van Erp, T. G. M., Sämann, P. G., Frodl, T., Jahanshad, N., Loehrer, E., Tiemeier, H., Hofman, A., Niessen, W. J., Vernooij, M. W., Ikram, M. A., Wittfeld, K., Grabe, H. J., Block, A., Hegenscheid, K., Völzke, H., Hoehn, D., Czisch, M., … Hibar, D. P. (2016). Subcortical brain alterations in major depressive disorder: Findings from the ENIGMA Major Depressive Disorder working group. Molecular Psychiatry, 21(6), 806–812. 10.1038/mp.2015.69

(0) Song, X. M., Hu, X.-W., Li, Z., Gao, Y., Ju, X., Liu, D.-Y., Wang, Q.-N., Xue, C., Cai, Y.-C., Bai, R., Tan, Z.-L., & Northoff, G. (2021). Reduction of higher-order occipital GABA and impaired visual perception in acute major depressive disorder. Molecular Psychiatry, 26(11), 6747–6755. 10.1038/s41380-021-01090-5

(1) Steffens, D. C., Wang, L., Manning, K. J., & Pearlson, G. D. (2017). Negative affectivity, aging, and depression: Results from the Neurobiology of Late-life Depression (NBOLD) study. The American Journal of Geriatric PsychiatrylJ: Official Journal of the American Association for Geriatric Psychiatry, 25(10), 1135–1149. 10.1016/j.jagp.2017.03.017

(2) Sudlow, C., Gallacher, J., Allen, N., Beral, V., Burton, P., Danesh, J., Downey, P., Elliott, P., Green, J., Landray, M., Liu, B., Matthews, P., Ong, G., Pell, J., Silman, A., Young, A., Sprosen, T., Peakman, T., & Collins, R. (2015). UK Biobank: An Open Access Resource for Identifying the Causes of a Wide Range of Complex Diseases of Middle and Old Age. PLOS Medicine, 12(3), e1001779. 10.1371/journal.pmed.1001779

(3) Tozzi, L., Garczarek, L., Janowitz, D., Stein, D. J., Wittfeld, K., Dobrowolny, H., Lagopoulos, J., Hatton, S. N., Hickie, I. B., Carballedo, A., Brooks, S. J., Vuletic, D., Uhlmann, A., Veer, I. M., Walter, H., Bülow, R., Völzke, H., Klinger-König, J., Schnell, K., … ‘for the ENIGMA-MDD Consortium.’ (2020). Interactive impact of childhood maltreatment, depression, and age on cortical brain structure: Mega-analytic findings from a large multi-site cohort. Psychological Medicine, 50(6), 1020–1031. 10.1017/S003329171900093X

(4) Wen, J., Fu, C. H. Y., Tosun, D., Veturi, Y., Yang, Z., Abdulkadir, A., Mamourian, E., Srinivasan, D., Skampardoni, I., Singh, A., Nawani, H., Bao, J., Erus, G., Shou, H., Habes, M., Doshi, J., Varol, E., Mackin, R. S., Sotiras, A., … iSTAGING consortium, A., Biocard, and BLSA. (2022). Characterizing Heterogeneity in Neuroimaging, Cognition, Clinical Symptoms, and Genetics Among Patients With Late-Life Depression. JAMA Psychiatry, 79(5), 464–474. 10.1001/jamapsychiatry.2022.0020

(5) Williams, L. M. (2016). Precision psychiatry: A neural circuit taxonomy for depression and anxiety. The Lancet. Psychiatry, 3(5), 472–480. 10.1016/S2215-0366(15)00579-9

(6) Winter, N. R., Leenings, R., Ernsting, J., Sarink, K., Fisch, L., Emden, D., Blanke, J., Goltermann, J., Opel, N., Barkhau, C., Meinert, S., Dohm, K., Repple, J., Mauritz, M., Gruber, M., Leehr, E. J., Grotegerd, D., Redlich, R., Jansen, A., … Hahn, T. (2022). Quantifying Deviations of Brain Structure and Function in Major Depressive Disorder Across Neuroimaging Modalities. JAMA Psychiatry, 79(9), 879–888. 10.1001/jamapsychiatry.2022.1780

(7) Wu, F., Lu, Q., Kong, Y., & Zhang, Z. (2023). A Comprehensive Overview of the Role of Visual Cortex Malfunction in Depressive Disorders: Opportunities and Challenges. Neuroscience Bulletin, 39(9), 1426. 10.1007/s12264-023-01052-7

58. Wüthrich, F., Lefebvre, S., Mittal, V. A., Shankman, S. A., Alexander, N., Brosch, K., Flinkenflügel, K., Goltermann, J., Grotegerd, D., Hahn, T., Jamalabadi, H., Jansen, A., Leehr, E. J., Meinert, S., Nenadić, I., Nitsch, R., Stein, F., Straube, B., Teutenberg, L., … Walther, S. (2023). The neural signature of psychomotor disturbance in depression. Molecular Psychiatry, 1–10. 10.1038/s41380-023-02327-1

(9) Xia, M., Si, T., Sun, X., Ma, Q., Liu, B., Wang, L., Meng, J., Chang, M., Huang, X., Chen, Z., Tang, Y., Xu, K., Gong, Q., Wang, F., Qiu, J., Xie, P., Li, L., He, Y., & DIDA-Major Depressive Disorder Working Group. (2019). Reproducibility of functional brain alterations in major depressive disorder: Evidence from a multisite resting-state functional MRI study with 1,434 individuals. NeuroImage, 189, 700–714. 10.1016/j.neuroimage.2019.01.074

60. Xiong, G., Dong, D., Cheng, C., Jiang, Y., Sun, X., He, J., Li, C., Gao, Y., Zhong, X., Zhao, H., Wang, X., & Yao, S. (2021). Potential structural trait markers of depression in the form of alterations in the structures of subcortical nuclei and structural covariance network properties. NeuroImage: Clinical, 32, 102871. 10.1016/j.nicl.2021.102871

61. Xu, Y., Wang, Y., Hu, N., Yang, L., Yu, Z., Han, L., Xu, Q., Zhou, J., Chen, J., Mao, H., & Pan, Y. (2022). Intrinsic Organization of Occipital Hubs Predicts Depression: A Resting-State fNIRS Study. Brain Sciences, 12(11), 1562. 10.3390/brainsci12111562

62. Yan, C.-G., Chen, X., Li, L., Castellanos, F. X., Bai, T.-J., Bo, Q.-J., Cao, J., Chen, G.-M., Chen, N.-X., Chen, W., Cheng, C., Cheng, Y.-Q., Cui, X.-L., Duan, J., Fang, Y.-R., Gong, Q.-Y., Guo, W.-B., Hou, Z.-H., Hu, L., … Zang, Y.-F. (2019). Reduced default mode network functional connectivity in patients with recurrent major depressive disorder. Proceedings of the National Academy of Sciences, 116(18), 9078–9083. 10.1073/pnas.1900390116

63. Yan, M., He, Y., Cui, X., Liu, F., Li, H., Huang, R., Tang, Y., Chen, J., Zhao, J., Xie, G., & Guo, W. (2021). Disrupted Regional Homogeneity in Melancholic and Non-melancholic Major Depressive Disorder at Rest. Frontiers in Psychiatry, 12, 618805. 10.3389/fpsyt.2021.618805

64. Yang, X., Kumar, P., Nickerson, L. D., Du, Y., Wang, M., Chen, Y., Li, T., Pizzagalli, D. A., & Ma, X. (2021). Identifying Subgroups of Major Depressive Disorder Using Brain Structural Covariance Networks and Mapping of Associated Clinical and Cognitive Variables. Biological Psychiatry Global Open Science, 1(2), 135–145. 10.1016/j.bpsgos.2021.04.006

65. Yoshida, K., Shimizu, Y., Yoshimoto, J., Takamura, M., Okada, G., Okamoto, Y., Yamawaki, S., & Doya, K. (2017). Prediction of clinical depression scores and detection of changes in whole-brain using resting-state functional MRI data with partial least squares regression. PloS One, 12(7), e0179638. 10.1371/journal.pone.0179638

66. Young, I. M., Dadario, N. B., Tanglay, O., Chen, E., Cook, B., Taylor, H. M., Crawford, L., Yeung, J. T., Nicholas, P. J., Doyen, S., & Sughrue, M. E. (2023). Connectivity model of the anatomic substrates and network abnormalities in major depressive disorder: A coordinate meta-analysis of resting-state functional connectivity. Journal of Affective Disorders Reports, 11, 100478. 10.1016/j.jadr.2023.100478

67. Zeng, L.-L., Shen, H., Liu, L., Wang, L., Li, B., Fang, P., Zhou, Z., Li, Y., & Hu, D. (2012). Identifying major depression using whole-brain functional connectivity: A multivariate pattern analysis. Brain, 135(5), 1498–1507. 10.1093/brain/aws059

68. Zhang, W., & Bijsterbosch, J. (2023). Dissociating Functional Brain Markers of State versus Trait Depression. 10.17605/OSF.IO/TXJMD

69. Zhao, Y., Yang, L., Sahakian, B. J., Langley, C., Zhang, W., Kuo, K., Li, Z., Gan, Y., Li, Y., Zhao, Y., Yu, J., Feng, J., & Cheng, W. (2023). The brain structure, immunometabolic and genetic mechanisms underlying the association between lifestyle and depression. Nature Mental Health, 1(10), Article 10. 10.1038/s44220-023-00120-1

70. Zhu, X., Yuan, F., Zhou, G., Nie, J., Wang, D., Hu, P., Ouyang, L., Kong, L., & Liao, W. (2021). Cross-network interaction for diagnosis of major depressive disorder based on resting state functional connectivity. Brain Imaging and Behavior, 15(3), 1279–1289. 10.1007/s11682-020-00326-2

71. Zhukovsky, P., Anderson, J. A. E., Coughlan, G., Mulsant, B. H., Cipriani, A., & Voineskos, A. N. (2021). Coordinate-Based Network Mapping of Brain Structure in Major Depressive Disorder in Younger and Older Adults: A Systematic Review and Meta-Analysis. American Journal of Psychiatry, 178(12), 1119–1128. 10.1176/appi.ajp.2021.21010088

