## Supplemental for "Higher amplitudes of visual networks are associated with trait-but not state-depression"

**Supplemental Materials**

**Participants**

To define participants in the state depression group, we applied a cut-off score of 𝜟RDS≥3 to capture significant changes in depression symptoms while maximizing the sample size. Varying the 𝜟RDS cut-off score (ranging from 1 to 6) resulted in different sample sizes and correlations between the measures used to define state and trait experiences of depression (***Table S1***).

**Table S1. Sample size of state depression group conditioned on different cut-off thresholds.**

| **Threshold** | **Representative range** ^a^ | **Resulting sample size** | **Correlation 𝜟RDS - RDS_baseline_** ^b^ |
| --- | --- | --- | --- |
| 𝜟RDS≥1 | 8% | 2,242 | 0.17 |
| 𝜟RDS≥2 | 17% | 847 | 0.11 |
| **𝜟RDS≥3** | **25%** | **311** | **0.06** |
| 𝜟RDS≥4 | 33% | 128 | 0.04 |
| 𝜟RDS≥5 | 42% | 59 | -0.13 |
| 𝜟RDS≥6 | 50% | 33 | -0.17 |

𝜟RDS = change scores in RDS between two scans (i.e., scan2 minus scan1)

^a^ The represented percentage of the full RDS scale (i.e., 16 points) for each chosen 𝜟RDS cut-off score.

^b^ The correlation between baseline RDS, used to define trait depression, and changes in RDS (i.e., 𝜟RDS), used to define state depression, varied across different 𝜟RDS cut-off scores. The selected cut-off score (in shade) exhibited a reduced correlation, with a substantial sample size.

**Group Matching**

The control participants were identified separately for each of the depression groups via a group matching method, which aimed to minimize the confounding effects of a set of covariates on potential group differences of interest. We considered several commonly used matching approaches that incorporated propensity score, including the nearest neighbor matching (Rosenbaum & Rubin, 1983, 1984), exact matching (Ho et al., 2007) (i.e., on sex), optimal matching (Gu & Rosenbaum, 1993; Rosenbaum, 1989), and genetic matching (Diamond & Sekhon, 2013; Tsai & Peace, 2011). The chosen propensity score-based *optimal matching* method (PS-Opt) exhibited the best performance compared to nearest neighbor (PS-NN), exact matching (PS-Ext) and genetic matching (PS-Gen; ***Table S3***), and the variables these groups were matched on showed no significant group differences (***Table S4***).

**Table S3. Diagnostics for matching methods under study.**

|  | **State (N=311)** | | | **Trait (N=265)** | | |
| --- | --- | --- | --- | --- | --- | --- |
| **Method** | **SMD** | **Var. Ratio** | **eCDFs** | **SMD** | **Var. Ratio** | **eCDFs** |
| **Pre-matching** | 1.1941 | 15.9838 | 0.36 | 0.8346 | 2.1102 | 0.2640 |
| **PS-NN** | 0.8380 | 12.2447 | 0.0728 | 0.0245 | 1.0674 | 0.0019 |
| **PS-Ext (sex)** | 0.8436 | 12.1958 | 0.0753 | 0.0357 | 1.0913 | 0.0032 |
| **PS-Opt** | **0.8377** | **12.2626** | **0.0728** | **0.0095** | **1. 0315** | **0.0011** |
| **PS-Gen** | 0.8768 | 11.0043 | 0.0970 | 0.0424 | 1.09218 | 0.0076 |

SMD = Standardized Mean Difference

Var. Ratio = Variance Ratio

eCDFs = Empirical Cumulative Distribution Functions

**Table S4. Mean values of matched variables after Optimal Matching.**

|  | **State-Control Pair** | | | **Trait-Control Pair** | | |
| --- | --- | --- | --- | --- | --- | --- |
|  | **State Depression** | **Control** | **Stats** ^a^ | **Trait Depression** | **Control** | **Stats** ^a^ |
| Sex-F ^b^ | 0.63 | 0.70 | 3.18 | 0.63 | 0.68 | 1.20 |
| Sex-M ^b^ | 0.37 | 0.30 |  | 0.37 | 0.33 |  |
| Age | 59.95 | 60.22 | 0.49 | 61.01 | 60.81 | -0.36 |
| Head Motion ^c^ | 0.12 | 0.12 | -0.96 | 0.12 | 0.11 | -0.95 |
| Site A ^c^ | 0.54 | 0.52 | 3.31 | 0.63 | 0.60 | 0.33 |
| Site B ^c^ | 0.05 | 0.03 |  | 0.10 | 0.11 |  |
| Site C ^c^ | 0.41 | 0.45 |  | 0.27 | 0.28 |  |
| Alcohol Intake | 1.88 | 1.92 | 1.19 | 1.86 | 1.87 | 0.22 |

^a^ Chi-square tests were conducted to examine between-group differences in categorical variables (i.e., sex, site), and paired t-tests to examine numerical variables.

^a^ Sex is coded as a categorical variable, including Sex-F (female) and Sex-M (male).

^c^ Variables relevant for brain measures including in-scanner head motion, and imaging acquisition site A-B.

Notably, control participants had minimal depression symptoms, with RDS ≤ 5 across all time points. This chosen cut-off indicates that at most one out of the four questions in the RDS scale was answered by ‘several days’, whereas all other questions by ‘not at all’, suggesting the minimal degree of depression symptomatology across all three timepoints. Additionally, control participants on antidepressants at any time point of assessments were excluded. We utilized information from the UKB variable (ID 20003, ‘Treatment/medication code’), obtained through a verbal interview with the question ‘In the touch screen you said you are taking regular prescription medications. Can you now tell me what these are?’. The name and corresponding data coding for each antidepressant medication in the UKB database are summarized in the list below:

*Antidepressants List (Index, Name, UKB data coding)*

1. allegron 10mg tablet, 1140867820
2. amitriptyline hydrochloride+perphenazine 10mg/2mg tablet, 1140867948
3. amitriptyline, 1140879616
4. amitriptyline+chlordiazepoxide 12.5mg/5mg capsule, 1140867938
5. anafranil10mg capsule, 1140867690
6. cipralex 5mg tablet, 1141190158
7. cipramil 10mg tablet, 1141151946
8. citalopram, 1140921600
9. clomipramine, 1140879620
10. cymbalta 30mg gastro-resistant capsule, 1141201834
11. depixol 3mg tablet, 1140867152
12. dosulepin, 1140909806
13. dothiepin, 1140879628
14. doxepin, 1140867640
15. duloxetine, 1141200564
16. edronax 4mg tablet, 1141151982
17. efexor 37.5mg tablet, 1140916288
18. escitalopram, 1141180212
19. faverin 50mg tablet, 1140867860
20. fluanxol 500micrograms tablet, 1140867952
21. fluoxetine, 1140879540
22. flupenthixol, 1140867150
23. flupentixol, 1140909800
24. fluphenazine hydrochloride+nortriptyline 1.5mg/30mg tablet, 1140867940
25. fluvoxamine, 1140879544
26. imipramine, 1140879630
27. isocarboxazid, 1140867856
28. lofepramine, 1140867726
29. lustral 50mg tablet, 1140867884
30. manerix 150mg tablet, 1140867922
31. maoi - tranylcypromine, 1140910820
32. mianserin, 1140879556
33. mirtazapine, 1141152732
34. moclobemide, 1140867920
35. molipaxin 50mg capsule, 1140882244
36. nardil 15mg tablet, 1140867852
37. nortriptyline, 1140867818
38. oxactin 20mg capsule, 1141174756
39. parnate 10mg tablet, 1140867916
40. paroxetine, 1140867888
41. phenelzine, 1140867850
42. prothiaden 25mg capsule, 1140867624
43. prozac 20mg capsule, 1140867876
44. reboxetine, 1141151978
45. seroxat 20mg tablet, 1140882236
46. sertraline, 1140867878
47. st john's wort/hypericum [ctsu], 1201
48. sinequan 10mg capsule, 1140882312
49. surmontil 10mg tablet, 1140867758
50. tofranil 10mg tablet, 1140867712
51. tranylcypromine, 1140867914
52. tranylcypromine+trifluoperazine 10mg/1mg tablet, 1140867944
53. trazodone, 1140879634
54. trimipramine, 1140867756
55. triptafen tablet, 1140867934
56. tryptophan product, 1140867960
57. venlafaxine, 1140916282
58. yentreve 20mg gastro-resistant capsule, 1141200570
59. zispin 30mg tablet, 1141152736

**fMRI Data Decomposition**

In the main analysis, the resting-state fMRI data were decomposed into 15 probabilistic modes or brain networks using the PRObabilistic Functional Modes (PROFUMO). PROFUMO is a hierarchical Bayesian approach that can decompose resting-state neuroimaging data into a set of modes or resting-state networks ^38,49^. Each estimated functional mode is described by a spatial map, network matrix (i.e., connectivity matrix – estimated as the partial correlation between mode time-courses), and mode amplitude. PROFUMO offers several advantages over alternative approaches, such as Independent Component Analysis (Beckmann et al., 2005) in conjunction with Dual Regression (Nickerson et al., 2017). Unlike these alternatives, PROFUMO circumvents the spatial independence constraint, allowing it to effectively capture overlapping network structures (Bijsterbosch et al., 2019). Moreover, PROFUMO employs an iterative optimization process for group and scan/subject estimates, leading to heightened sensitivity in discerning individual-specific network organization (Bijsterbosch et al., 2018; Harrison et al., 2020).

To identify spurious modes, characterized by either being consistently empty across participants/timepoints (indicated by low mode amplitudes and/or spatial weights) or containing noise (e.g., activity in the circle of Willis), we conducted visual inspections of group-level spatial maps for each PROFUMO mode. Two spurious modes were excluded from the total of 15 decomposed modes, resulting in n=13 meaningful brain networks for statistical analyses. These functional modes largely resemble canonical resting-state networks, including the default mode, central executive, salience, visual, and motor networks (***Figure S1***).

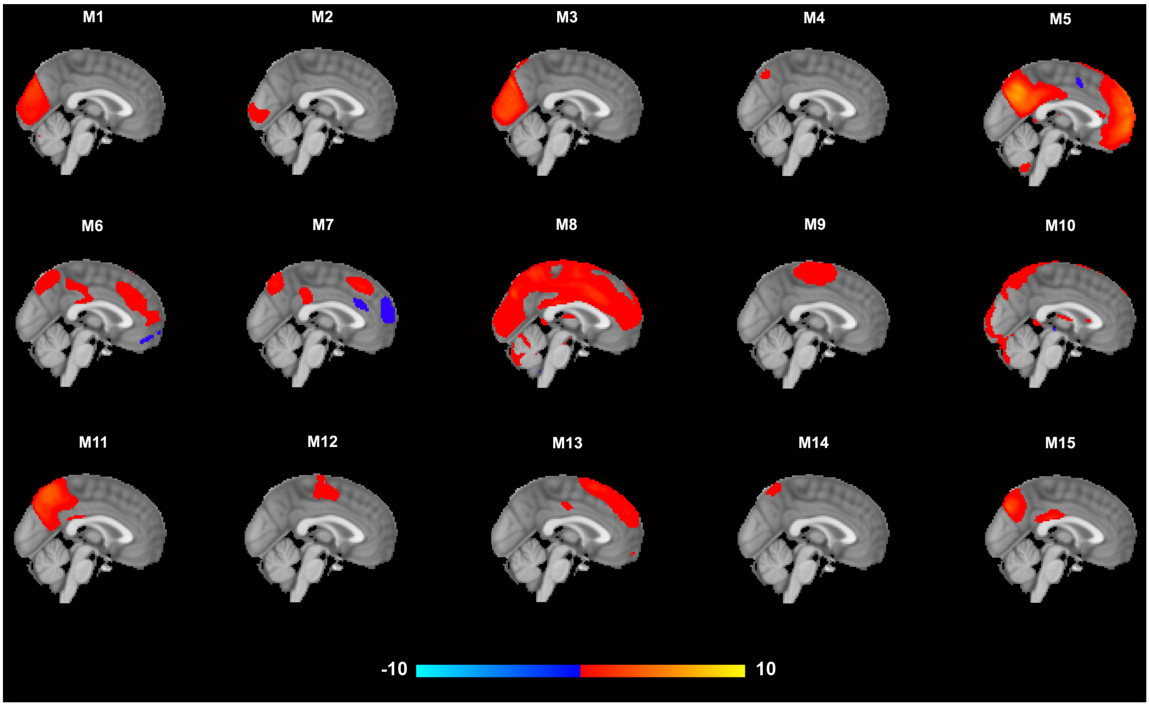

**Figure S1.** The decomposed 15 functional modes/networks, including two spurious modes (M8 and M10) that were excluded from statistical analyses.

To replicate the significant findings observed in the 15 modes, we further decomposed the data with 10 and 20 dimensions, separately. Similar resemblances with canonical networks can be observed in these decompositions with varying dimension decompositions, with some of the resulting networks in one decomposition merging into one or splitting into two or more modes in another decomposition (***Figure S2***; ***Figure S3***).

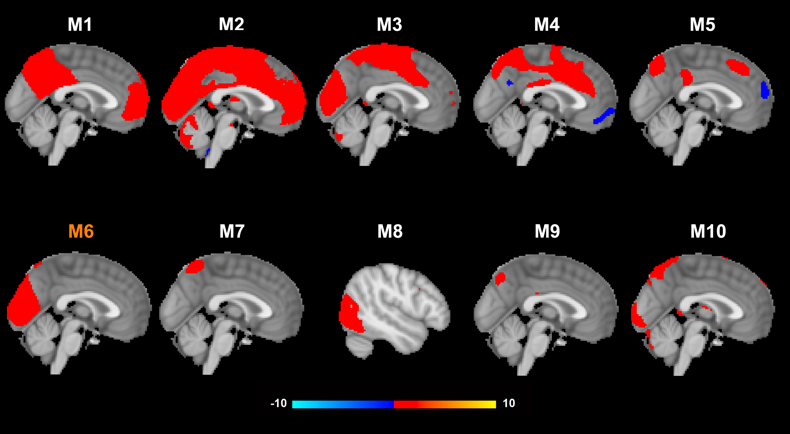

**Figure S2.** The decomposed 10 functional modes/networks, including two spurious modes (M2 and M10). Mode 6 (M6) from this decomposition showed the highest Cosine similarity with mode 2 and mode 3 from 15-dimension decomposition, the amplitude of which was found higher in individuals with trait experience of depression in contrast to control participants.

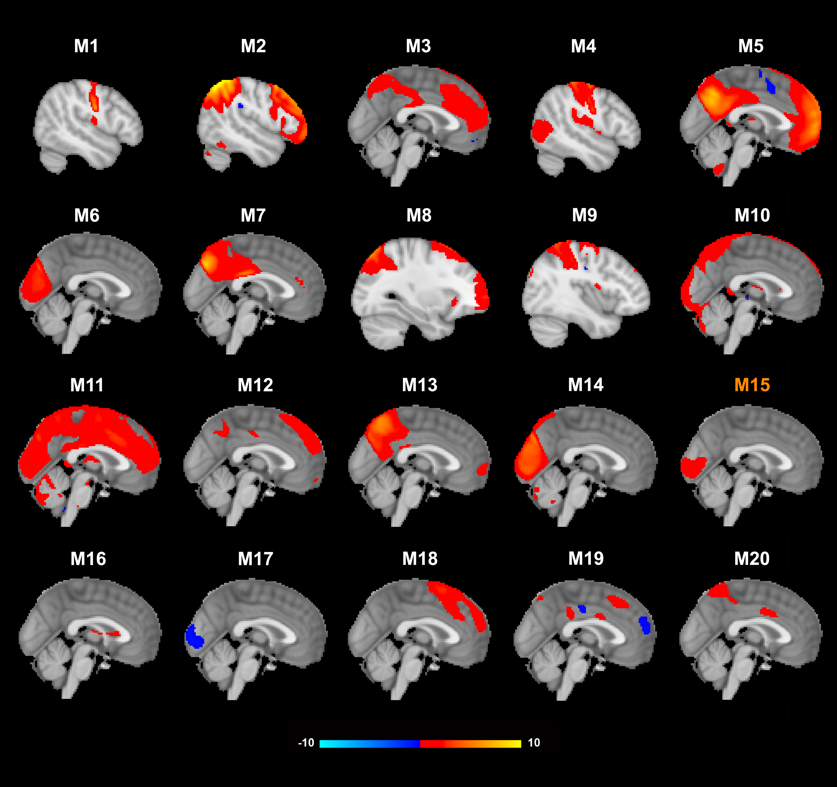

**Figure S3.** The decomposed 20 functional modes/networks, including three spurious modes. (M10, M11 and M16). Mode 15 (M15) from this decomposition showed the highest Cosine similarity with mode 2 from 15-dimension decomposition, the amplitude of which was found higher in individuals with trait experience of depression in contrast to control participants.

**Supplemental Results**

To validate the robustness of our main findings, which indicated that individuals with a trait experience of depression exhibited higher amplitude in visual networks compared to control participants, we repeated these analyses using 10 and 20 decomposition dimensionalities, which resulted in 10 and 20 brain networks (rather than 15), respectively.

We first identified the “best-matched” modes from 10- and 20-dimension decompositions for the target modes showing the significant between-group difference in amplitudes. Cosine similarity coefficients were used and the identified modes that exhibited the most similar spatial distributions to the target ones from the 15-dimension decomposition (***Figure S2-3***).

Consistently, we observed the same pattern as in our main findings: higher amplitude in the visual networks was noted in participants with trait depression compared to controls (β’s > 0.02, p’s < 0.03), even when brain networks were identified with different decomposition dimensionalities. Intriguingly, the two visual networks from the 15-mode decomposition appeared to merge into one network (i.e., mode 6) in the 10-mode decomposition, while the same visual networks maintained their spatial distribution in only one network (i.e., mode 15) in the 20-mode decomposition (***Figure S2-3***; ***Table S5***). These corresponding networks were identified based on their Cosine similarity with the target modes (see full similarity matrices in ***Figure S4***).

**Table S5. Replicated results of higher visual network amplitude.**

|  | **β** | **P-value** ^a^ |
| --- | --- | --- |
| Visual Network from 10-Mode Decomposition | 0.05 | <0.01 |
| Visual Network from 20-Mode Decomposition | 0.02 | <0.05 |

^a^ Uncorrected p-values were reported for these replication analyses that only tested the significant findings from the main analyses.

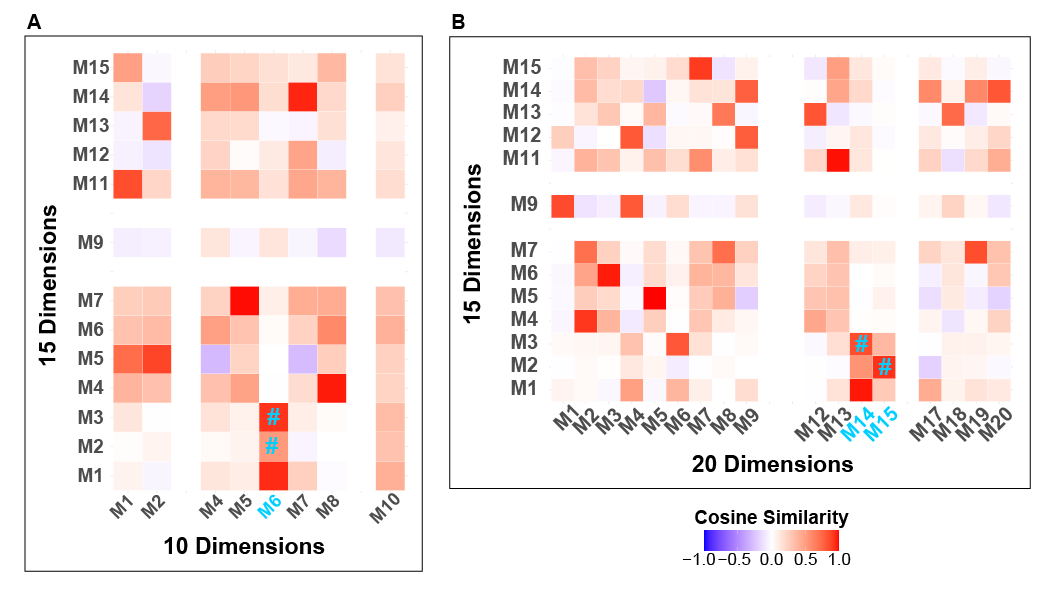

**Figure S4.** Similarity matrices between each pair of modes from 15- and 10-dimension decompositions (panel A), and between 15- and 20-dimension decompositions (panel B). M1-M20 indicates mode1-mode20, in different decompositions. The spurious modes from each decomposition were removed from the matrices (i.e., empty rows and columns) and mode 6 (M6) from 10-dimension decomposition showed the highest Cosine similarity to mode 2 and mode 3 from 15-dimension decomposition (i.e., two visual networks), as highlighted in blue and with *#* symbol, whereas mode15 (M15) and mode 14 (M14) from 20-dimension decomposition “best matched” mode 2 and 3 from 15-dimension decomposition, respectively.

**Reference:**

Beckmann, C. F., DeLuca, M., Devlin, J. T., & Smith, S. M. (2005). Investigations into resting-state connectivity using independent component analysis. *Philosophical Transactions of the Royal Society B: Biological Sciences*, *360*(1457), 1001–1013. https://doi.org/10.1098/rstb.2005.1634

Bijsterbosch, J. D., Beckmann, C. F., Woolrich, M. W., Smith, S. M., & Harrison, S. J. (2019). The relationship between spatial configuration and functional connectivity of brain regions revisited. *eLife*, *8*, e44890. https://doi.org/10.7554/eLife.44890

Bijsterbosch, J. D., Woolrich, M. W., Glasser, M. F., Robinson, E. C., Beckmann, C. F., Van Essen, D. C., Harrison, S. J., & Smith, S. M. (2018). The relationship between spatial configuration and functional connectivity of brain regions. *eLife*, *7*, e32992. https://doi.org/10.7554/eLife.32992

Diamond, A., & Sekhon, J. S. (2013). Genetic Matching for Estimating Causal Effects: A General Multivariate Matching Method for Achieving Balance in Observational Studies. *The Review of Economics and Statistics*, *95*(3), 932–945.

Gu, X. S., & Rosenbaum, P. R. (1993). Comparison of Multivariate Matching Methods: Structures, Distances, and Algorithms. *Journal of Computational and Graphical Statistics*, *2*(4), 405–420. https://doi.org/10.1080/10618600.1993.10474623

Harrison, S. J., Bijsterbosch, J. D., Segerdahl, A. R., Fitzgibbon, S. P., Farahibozorg, S.-R., Duff, E. P., Smith, S. M., & Woolrich, M. W. (2020). Modelling subject variability in the spatial and temporal characteristics of functional modes. *NeuroImage*, *222*, 117226. https://doi.org/10.1016/j.neuroimage.2020.117226

Ho, D. E., Imai, K., King, G., & Stuart, E. A. (2007). Matching as Nonparametric Preprocessing for Reducing Model Dependence in Parametric Causal Inference. *Political Analysis*, *15*(3), 199–236. https://doi.org/10.1093/pan/mpl013

Nickerson, L. D., Smith, S. M., Öngür, D., & Beckmann, C. F. (2017). Using Dual Regression to Investigate Network Shape and Amplitude in Functional Connectivity Analyses. *Frontiers in Neuroscience*, *11*, 115. https://doi.org/10.3389/fnins.2017.00115

Rosenbaum, P. R. (1989). Optimal Matching for Observational Studies. *Journal of the American Statistical Association*, *84*(408), 1024–1032. https://doi.org/10.1080/01621459.1989.10478868

Rosenbaum, P. R., & Rubin, D. B. (1983). The central role of the propensity score in observational studies for causal effects. *Biometrika*, *70*(1), 41–55. https://doi.org/10.1093/biomet/70.1.41

Rosenbaum, P. R., & Rubin, D. B. (1984). Reducing Bias in Observational Studies Using Subclassification on the Propensity Score. *Journal of the American Statistical Association*, *79*(387), 516–524. https://doi.org/10.1080/01621459.1984.10478078

Tsai, K.-T., & Peace, K. (2011). Genetic Matching: A Better Algorithm for Adjusting Covariate Imbalance for Statistical Data Analysis and Modeling. *Proceedings of the International Conference on Bioinformatics and Computational Biology*. https://digitalcommons.georgiasouthern.edu/biostat-facpubs/54
